# Functional characterization of the store-operated calcium entry pathway in naked mole-rat cells

**DOI:** 10.1101/2024.12.15.628530

**Authors:** Polina Drugachenok, Paulina Urriola-Muñoz, Lanhui Qui, Zhuang Zhuang Han, Ewan St J Smith, Taufiq Rahman

## Abstract

Naked mole-rats (NMRs, *Heterocephalus glaber*) are highly unusual rodents exhibiting remarkable adaptations to their subterranean habitat and resistance to developing various age-related diseases such as those related to abnormal cell proliferation or cancer, neurodegeneration and inflammation. In other rodents, as well as humans, a ubiquitous Ca^2+^ influx pathway, namely the store-operated Ca^2+^ entry (SOCE), has been implicated in all these diseases. SOCE is triggered by intracellular Ca^2+^ store depletion resulting in interaction of Stim proteins with Orai proteins, the putative homologs of which appear be present in the NMR genome, but no characterisation of SOCE in NMRs has yet been conducted. In this study, we provide the first functional and pharmacological characterization of SOCE in NMR cells using both excitable and non-excitable cells.

## INTRODUCTION

The store-operated Ca^2+^ entry (SOCE) represents a route through which Ca^2+^ from the extracellular fluid enters into the cytosol in response to depletion of intracellular Ca^2+^ stores, such as the endoplasmic reticulum (ER)[1]. Store depletion occurs physiologically upon stimulation of cell surface receptors that produce the second messenger - inositol 1, 4, 5-trisphosphate (IP_3_) in a phospholipase C (PLC)-dependent manner. Pharmacologically, SOCE can also be triggered either by inhibiting (e.g. with thapsigargin or cyclopiazonic acid) the SarcoEndoplasmic Reticulum Ca^2+^-ATPase (SERCA) pump, which allows uncompensated passive Ca^2+^ leak from the ER that eventually empties the store, or by using a low concentration (≤10μM) of ionomycin that mobilises Ca^2+^ preferentially from the intracellular stores, thereby depleting them.

Initially SOCE was thought to be an exclusive feature of the non-excitable cells, but more recent studies have confirmed that this particular Ca^2+^ influx pathway is present almost in all cells including the excitable ones, although its prominence and indispensability can vary depending on the nature of the cell[1-3]. In most eukaryotic cell types, SOCE is mediated through Ca^2+^ release-activated Ca^2+^ (CRAC) channels. The CRAC channels are highly Ca^2+^-selective and, rather unusually for ion channels, formed *in situ*. CRAC channel activation occurs following store depletion that induces interaction between clusters of the plasma membrane (PM)-resident Orai proteins and the ER-membrane localized STIM proteins at ER-PM junctions. Evidence continues to amass that CRAC-derived Ca^2+^ signals, besides their well-known ER-refilling housekeeping role, are often utilized for regulating specific cellular functions including gene transcription, secretion, modulation of enzyme activity and motility [1, 3]. Abnormal SOCE due to aberrant CRAC channel activity has been implicated in various diseases, including various autoimmune and inflammatory diseases, neurodegenerative diseases, chronic pain and cancer [4, 5].

The naked mole-rat (NMR, *Heterocephalus glaber*) is a long-lived, subterranean, and eusocial mammal, which exhibits numerous signs of healthy aging alongside a remarkable tolerance to hypoxia and unusual pain phenotypes, as well as notable resistance towards developing certain age-associated diseases, such as cancer, neurodegenerative conditions and cardiovascular ailments[6]. Across pain, cancer and neurodegeneration, the immune system plays an important role. Intriguingly, when compared to standard laboratory rodents, the NMR immune system manifests some unusual features, such as absence of natural killer cells, higher pro-inflammatory cytokine production by macrophages and a novel neutrophil subset[7]. It is well-established that CRAC channels mediating SOCE are critical for proper functioning of various immune cells [8], such that aberrant SOCE due to abnormal CRAC channel activity is involved in various chronic inflammatory diseases [4] to which NMRs are comparatively more resistant. In addition to the role of CRAC channels in regulating immune cell function, research also indicates that Stim and Orai proteins are expressed in dorsal root ganglion (DRG) sensory neurons and spinal cord dorsal horn neurons, thus suggesting that they play a role in nociceptive processing and pain[6, 9]. We therefore sought to functionally characterize SOCE and its underlying CRAC channel components in NMR cells to determine whether this pathway exhibits any unusual features that might potentially contribute to its unusual biology.

## MATERIALS AND METHODS

### Cell Culture

Immortalised fibroblasts from NMR skin (‘NMR SV40 fibroblasts’) were established in the Smith laboratory previously[10] and cultured as per published protocol. Briefly, cells were maintained in Dulbecco’s modified Eagle’s medium (DMEM) high glucose (Gibco™) supplemented with 15% (v/v) fetal bovine serum (Sigma-Aldrich™), non-essential amino acids, 1mM sodium pyruvate, 100 units/mL penicillin and 100□mg/mL streptomycin (Gibco™). Cells were incubated in a humidified 32°C incubator with 5% CO_2_ and 3% O_2_. NMR DRG neurons (‘NMR-DRG neurons’) were isolated and cultured as per published protocol [11]. NMR were sacrificed using a rising CO_2_ concentration followed by decapitation in accordance with the UK Animal (Scientific Procedures) Act 1986 Amendment Regulations 2012 under a Project License (PP5814995) granted to E. St. J. S. by the Home Office with additional approval by the University of Cambridge Animal Welfare Ethical Review Body. Before collecting the NMR-DRG neurons, sterile 35mm glass-bottomed petri dishes (MatTek, USA) were coated with laminin (20μg/mL) and incubated at 37°C for 1 hour. Once plated, NMR-DRG neurons were incubated in a humidified 37°C, 5% CO_2_ incubator overnight before Ca^2+^ imaging the following day. Jurkat T cells were cultured as per previously published protocol [12].

### Ca^2+^ Imaging

All Ca^2+^ imaging experiments involving the NMR-SV40 fibroblasts and NMR-DRG neurons were performed at room temperature (21°C-23°C) using Fura-2 based Ca^2+^ imaging as per previously published protocol[12]. Paired fluorescence images (340 and 380 nm excitation, 510 nm emission) were captured every 5s using a QIClick™ digital CCD camera (QImaging, BC, Canada) mounted on a Nikon Eclipse Ti-S Microscope with MetaFluor® (Molecular Devices, USA) Fluorescence Ratio Imaging Software.

### Immunocytochemistry and confocal microscopy

Jurkat T cells or NMR SV40 fibroblasts plated on PDL-coated glass coverslips were fixed using formaldehyde or methanol, respectively and permeabilised using Triton X-100 following published protocols[13, 14]. Fixed cells were treated overnight at 4°C with polyclonal rabbit FITC-conjugated antibodies (Cusabio®, Texas, US) raised against human Orai, hOrai1 (product code: CSB-PA846601LC01HU) and human Stim1, hStim1 (produce code: CSB-PA022829LC01HU) at a final concentration of 25μg/mL in phosphate buffered saline (PBS). After overnight incubation, coverslips were washed with PBS thrice and treated with 1μg/mL nuclear stain DAPI (Sigma-Aldrich) for 5 minutes at room temperature. The coverslips were then washed thrice with PBS and mounted onto a microscope slide using ProLong™ Gold Antifade Mountant (ThermoFisher Scientific, UK). Confocal images were acquired using a Leica SP5 confocal microscope equipped with a 63X oil immersion lens (Leica Microsystems, Wetzlar, Germany). The excitation wavelengths used were 405 nm for DAPI and 488 nm for FITC. Acquired images were processed using Leica Application Suite X (LAS X).

### Statistical analysis

Results were expressed as means ± SEM for n independent experiments done in 3 different days. Statistical comparisons of the mean values were performed in GraphPad Prism 9 (GraphPad software Inc., CA, USA) using one-way ANOVA followed by Tukey’s multiple comparisons test or Student’s t-test (unpaired, two-tailed); p<0.05 was considered to be significant.

## RESULTS AND DISCUSSION

### Preliminary Characterization of SOCE from NMR SV40 cells

We first wanted to determine the likelihood that the protein machinery underpinning SOCE or CRAC channels, namely Orai and Stim, are expressed by NMR cells. Therefore, we looked up the NMR genome database (http://www.naked-mole-rat.org/) as well as Uniprot (https://www.uniprot.org/) and identified amino acid sequences representing the putative NMR Orai and Stim isoforms, significant (>85%) similarities being shared with cognate sequences of mouse, rat and human Orai and Stim isoforms (**Supplementary Figs. S1-S5**).

Since Orai1 and Stim1 are the major isoforms that underlie native SOCE in most mammalian cells[3], we conducted immunostaining of immortalized NMR skin fibroblasts (NMR SV40 fibroblasts, developed previously in the Smith lab[10], using FITC-conjugated antibodies raised against human Stim1 (hStim1) and human Orai1 (hOrai1) protein, being able to detect both Stim1 and Orai1 (**Fig.1A**). The fidelities of these antibodies were assessed in Jurkat T cells (**Supplementary Fig.S6**) that are well-known to express hStim1 and hOrai1 isoforms and manifest robust SOCE[1, 3, 12].

**Figure 1.**
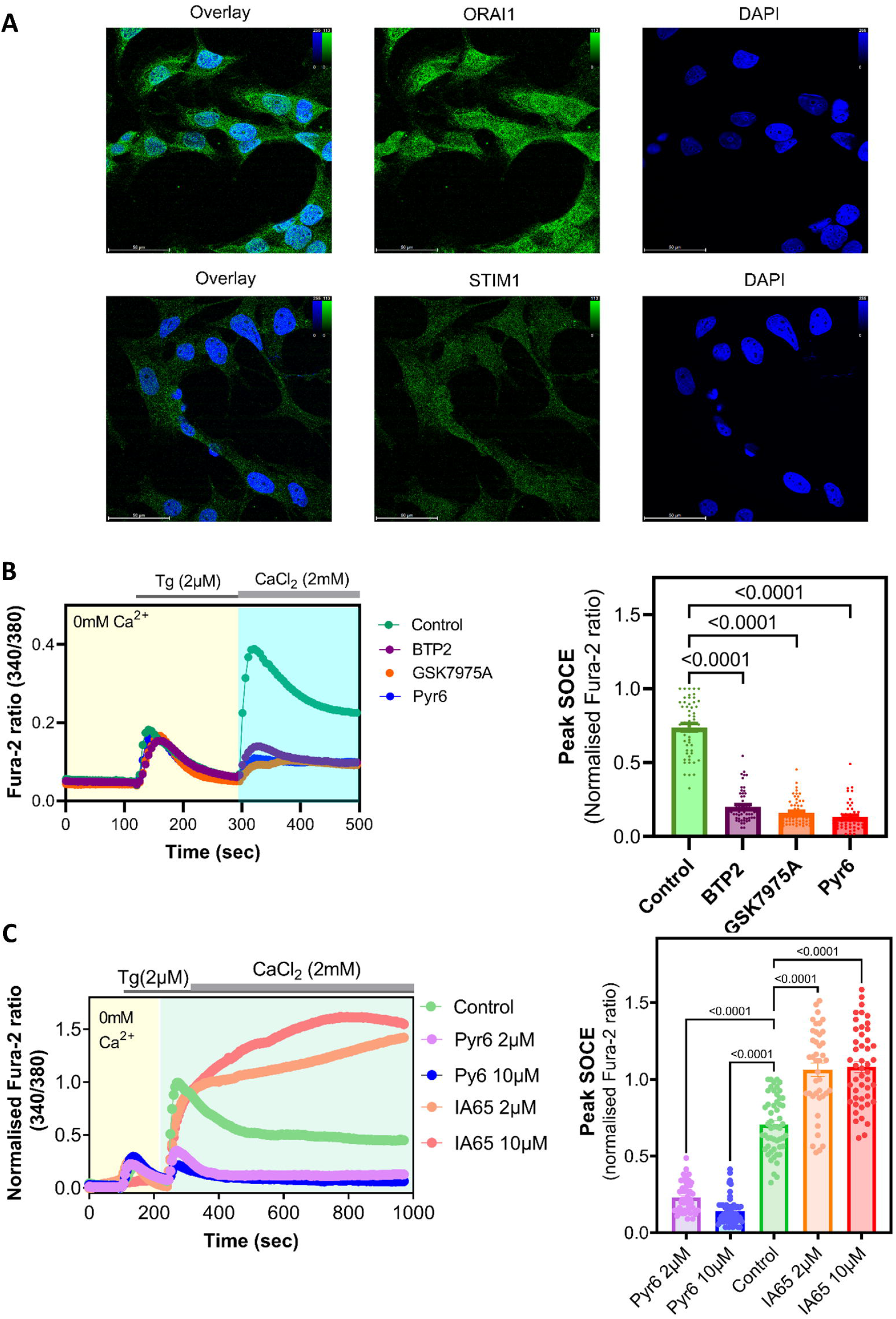
Preliminary characterization of SOCE and its underlying protein players in NMR SV40 fibroblasts. **A**) Exemplar confocal images showing expression of endogenous Orai and Stim proteins in NMR SV40 fibroblasts. The latter were fixed and stained with FITC-conjugated antibodies raised against hOrai1 and Stim1; nuclei were stained with DAPI and shown in blue. Scale bar, 50μm. **B**) Left, exemplar traces showing Ca^2+^ signals (indicated by Fura-2 fluorescence ratio) triggered by adding 2□μM of thapsigargin (Tg) to NMR SV40 fibroblastss with or without pre-treatment (10μM for 10min) with the indicated known SOCE/CRAC channel inhibitors (shown in different non-green colours). Right, summary data obtained from Ca^2+^ imaging experiments shown in left, showing peak Ca^2+^ entry (SOCE) observed in NMR SV40 fibroblasts under different conditions. **C**) Left, exemplar traces showing Tg-evoked Ca^2+^ signals to NMR SV40 fibroblastss with or without pre-treatment (2 or 10μM, for 10min) with Pyr6 (known SOCE/CRAC channel inhibitor) and IA65 (known SOCE/CRAC channel enhancer). Right, summary data obtained from Ca^2+^ imaging experiments shown in left, showing peak Ca^2+^ entry (SOCE) observed in NMR SV40 fibroblasts under different conditions. Data are mean ± sem from ≥90 cells (per each experimental condition) plated on 3 petri dishes, each imaged on different days. Averages from the experimental groups (i.e. different SOCE-inhibitor treated ones) were compared to that of the control group using one-way ANOVA followed by Tukey’s multiple comparisons test; p<0.05 was considered to be significant.

After confirming the presence of SOCE components at both genomic and protein levels, we next sought to functionally characterize SOCE in NMR SV40 fibroblasts. For this, we used Fura-2 based Ca^2+^ imaging as per published protocol[12], initially depleting internal stores with 2 μM thapsigargin (Tg) in the absence of extracellular Ca^2+^, followed by addition of Ca^2+^ (as 2 mM CaCl_2_). This protocol induced a robust Ca^2+^-entry rapidly after the addition of Ca^2+^ to the extracellular solution bathing the NMR SV40 fibroblasts (**Fig.1B**). The Ca^2+^ response was significantly attenuated when NMR SV40 fibroblasts were preincubated for 10-minutes with 10 μM BTP2, GSK-7975Aor Pyr-6, each being a widely-used blocker of SOCE/CRAC channels[2, 12] (**Fig.1B**). In a separate series of Ca^2+^ imaging experiments, we used the same protocol and recorded Tg-evoked Ca^2+^ entry in the NMR SV40 fibroblasts pretreated with either low (2 μM) or high (10 μM) concentration of Pyr6, or the recently-discovered enhancer of Orai1 namely IA65[15]. Compared to untreated cells, Tg-evoked Ca^2+^ entry in Pyr6 pretreated NMR SV40 fibroblasts was significantly attenuated, whereas pretreatment with IA65 treatment resulted in a significantly greater Ca^2+^ response (**Fig.1C**). Thus, these two well-characterized SOCE-selective pharmacological tools modulated NMR SOCE in a similar to way that the published literature shows that they do in other mammalian cell types [15]. We noticed that 2 μM Pyr6 sufficed at causing significant suppression of SOCE in NMR SV40 fibroblasts and that 10 μM did not produce any greater attenuation. Thus, in NMR SV40 fibroblasts we were able to detect clear, pharmacologically, i.e. Tg-evoked SOCE representing active CRAC channels that were sensitive to well-known pharmacological modulators of this pathway.

Next, we aimed to determine whether SOCE could be physiologically evoked in NMR SV40 fibroblasts, i.e through depleting the ER Ca^2+^ store by stimulating cells to activate PLC isoforms and thereby generate the Ca^2+^-mobilising second messenger IP_3_. Given no a priori knowledge was available regarding the presence of specific Gq-coupled receptors or receptor tyrosine kinases in NMR SV40 fibroblasts, we arbitrarily tested few agonists for cell surface receptors including ATP, endothelin-1 and histamine that are well-known to activate PLC and produce IP_3_ in many eukaryotic cell types[16]. However, none of these agonists produced any discernible Ca^2+^ signal in the NMR SV40 fibroblasts (results not shown). By contrast, angiotensin II (Ang II), an endogenously-produced octapeptide and a highly potent vasoconstrictor[17], was observed to mobilise Ca^2+^ in NMR SV40 fibroblasts and deplete Ca^2+^ stores, such that Ca^2+^ addition led to robust Ca^2+^ entry via SOCE, as indicated by inhibition by Pyr6 **Fig.2**. Thus, these NMR fibroblasts are likely to express AT_1_ receptors that are G_q_-coupled G protein-coupled receptors, stimulation of which is known to deplete ER Ca^2+^ stores and trigger SOCE[17, 18]. This result also suggests that NMRs, like other mammals, are able to employ SOCE to conduct various cellular functions, including the housekeeping store refilling purpose[1, 3].

**Figure 2.**
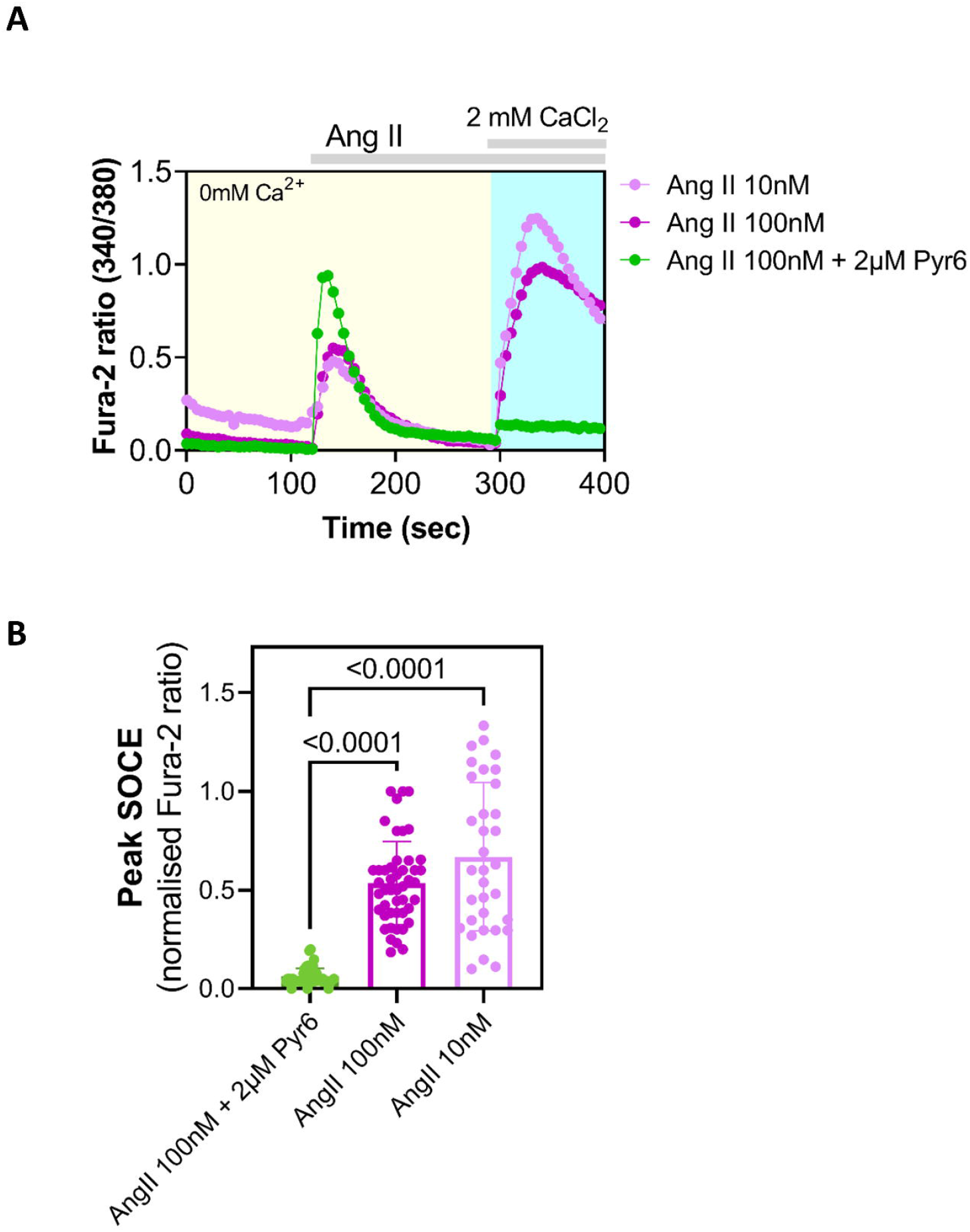
Physiologically-evoked SOCE in NMR SV40 fibroblasts. **A**) Exemplar traces showing Ca^2+^ signals (indicated by Fura-2 fluorescence ratio) triggered in NMR fibroblasts by adding indicated concentrations of Angiotensin II (Ang-II) alone or together with Pyr6 in Ca^2+^-free extracellular media followed by subsequent Ca^2+^ add back. **B**) Summary data obtained from Ca^2+^ imaging experiments shown in left, showing peak Ca^2+^ entry (SOCE) observed in NMR SV40 fibroblasts under different conditions. Data are mean ± sem from ≥75 cells (per each experimental condition) plated on 3 petri dishes, each imaged on different days. Averages from the experimental groups (i.e. different SOCE-inhibitor treated ones) were compared to that of the control group using one-way ANOVA followed by Tukey’s multiple comparisons test; p<0.05 was considered to be significant.

### Evaluating the effect of exposure to different oxygen levels on SOCE in NMR SV40 fibroblasts

It is well-documented that NMR fibroblasts thrive under lower O_2_ culturing conditions (3%) than cells from mouse or human, perhaps an adaptation to their subterranean environment and in agreement with their hypoxia resistance [10, 14, 19]. However, all Ca^2+^ imaging experiments, with or without pre-incubation with various SOCE modulators, were performed at atmospheric O_2_ levels and in HBSS (**Figs1B-C** and **Fig. 2**), which might not be optimal for culturing and maintaining the NMR SV40 fibroblasts. Although, NMR SV40 fibroblasts looked healthy during the time period in atmospheric O_2_ and responded well to our Ca^2+^ imaging protocol, we nevertheless sought to explore whether or not changes in the O_2_ level could affect SOCE in these cells. Accordingly, in subsequent experiments we performed all the pre-Ca^2+^ imaging preparations such as Fura-2AM loading, washing and de-esterification etc. in the NMR SV40 fibroblast maintained in optimal media (cRPMI) and optimal oxygen level (3% O_2_) and compared this to when procedures were conducted in sub-optimal media (HBSS) and optimal oxygen level (3% O_2_) and in sub-optimal media (HBBS) and sub-optimal/atmospheric oxygen level (21% O_2_). Regardless of the pre-Ca^2+^-imaging conditions, we measured SOCE in our usual recording conditions of room temperature and atmospheric O_2_ and in HBSS. When compared with one another, the peak Ca^2+^ entry via SOCE was comparable across different cohorts of NMR SV40 fibroblasts (**Fig.3**), thus suggesting that short-term changes in atmospheric O_2_ did not overtly compromise the function of NMR SV40 fibroblast SOCE.

**Figure 3.**
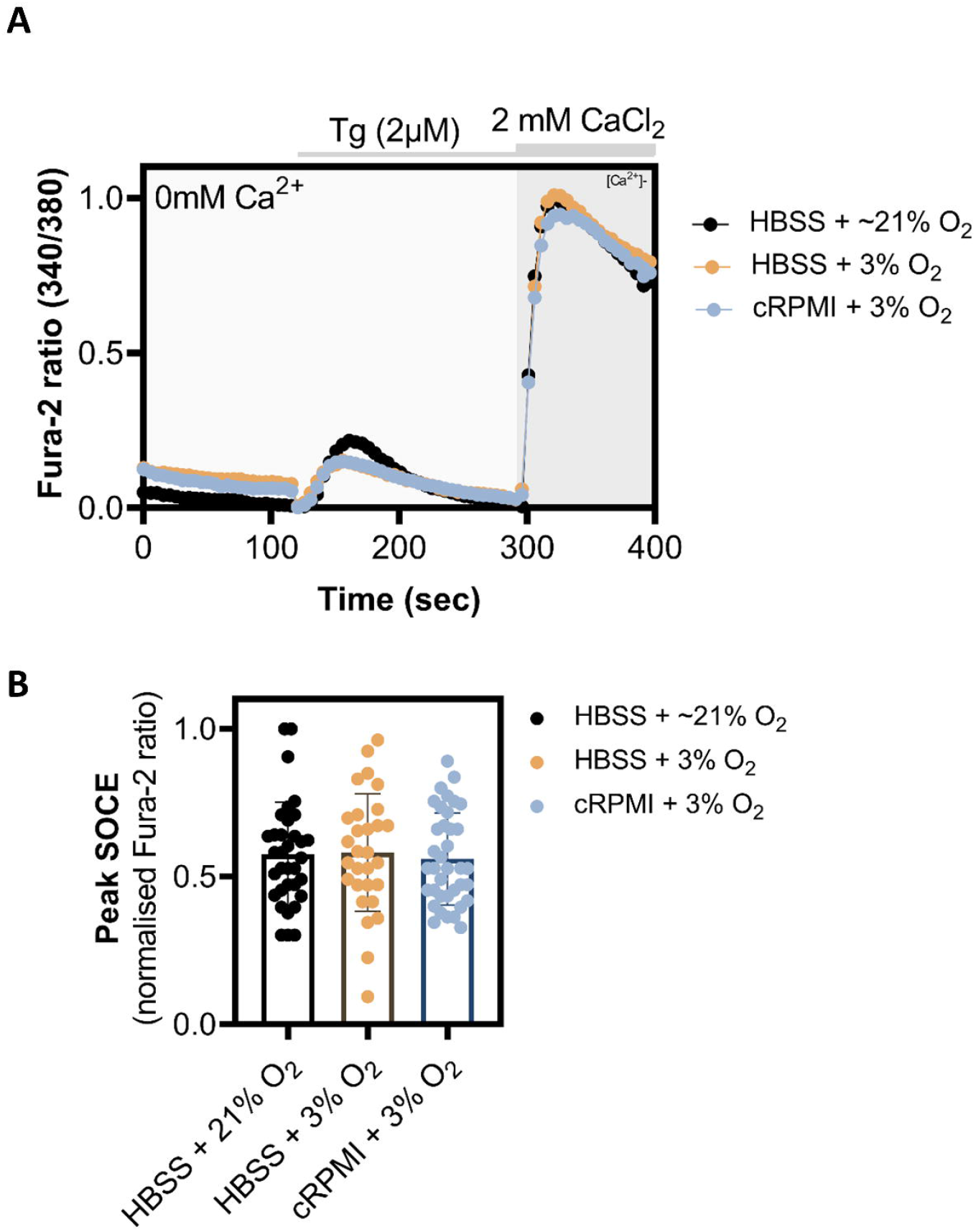
Evaluation of exposure to different oxygen level on SOCE in NMR SV40 fibroblasts. **A**) Exemplar Ca^2+^ imaging traces showing Tg-evoked SOCE (indicated by Fura-2 fluorescence ratio) in NMR SV40 fibroblasts for which loading of Fura-2AM dye and its subsequent de-esterification was carried out in suboptimal media and optimal O_2_ level (HBSS, 21% O_2_), suboptimal media and optimal O_2_ level (HBSS, 3% O_2_) and optimal media and optimal O_2_ level (cRPMI, 21% O_2_). **B**) Summary data obtained from Ca^2+^ imaging experiments shown in left, showing peak Ca^2+^ entry (SOCE) observed in NMR SV40 fibroblasts under different conditions. Data are mean ± sem from ≥75 cells (per each experimental condition) plated on 3 petri dishes, each imaged on different days. Averages from the experimental groups (i.e. different SOCE-inhibitor treated ones) were compared to that of the control group using one-way ANOVA followed by Tukey’s multiple comparisons test; p<0.05 was considered to be significant.

### Characterising SOCE in primary Naked Mole-Rat neurons

Characterisation of NMR SOCE in immortalized NMR fibroblasts does not necessarily serve as true representative of NMR SOCE and we therefore sought to characterise SOCE in NMR primary cells. Given the fact that SOCE has been well documented in mammalian DRG neurons[20] and our experience with these cells[11, 21], we decided to characterise SOCE in NMR DRG neurons. As in NMR SV40 fibroblasts, we were able to detect robust SOCE both pharmacologically (i.e. using Tg to passively deplete the ER Ca^2+^ store, **Fig. 4A**) as well as physiologically (using Ang II to deplete the ER Ca^2+^ store through generating IP_3_, **Fig. 4B**), SOCE being significantly suppressed in both cases when pretreated with the SOCE inhibitor Pyr6 (2 μM). At the end of each experiment, 50 mM KCl was added to depolarise the neurons and trigger Ca^2+^ entry via voltage-gated Ca^2+^ channels, only such cells, thus identified as neurons, were analysed for SOCE activity.

**Figure 4.**
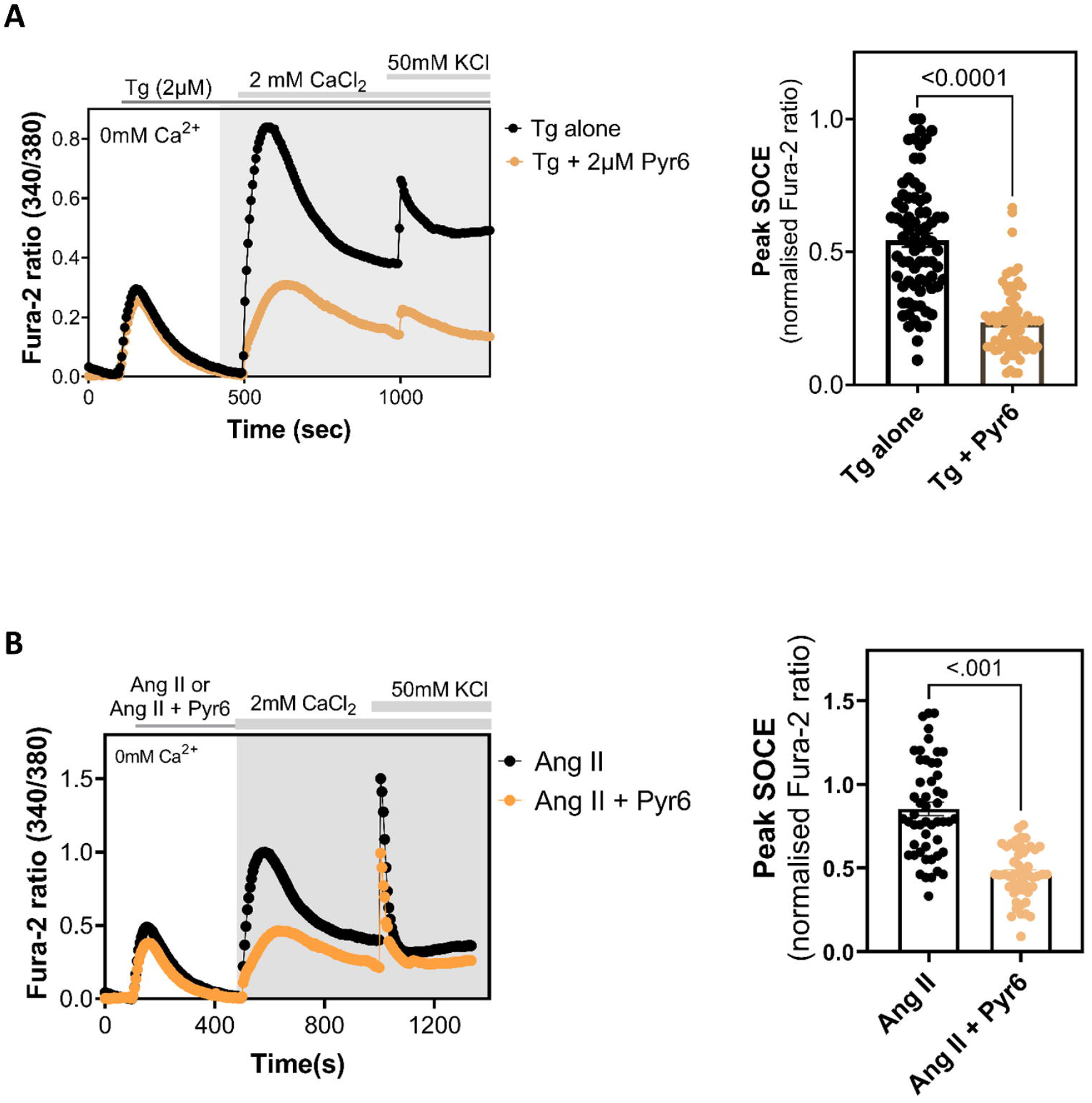
SOCE in DRG neurons isolated from the nake mole rats. **A**) Left, exemplar traces showing Ca^2+^ signals (indicated by Fura-2 fluorescence ratio) triggered in DRG neurones from NMR by adding Tg (2μM) alone or together with Pyr6 (2μM) in Ca^2+^-free extracellular media followed by subsequent Ca^2+^ add back. Right, summary data obtained from Ca^2+^ imaging experiments shown in left, showing peak Ca^2+^ entry (SOCE) observed in DRG neurones from NMR under different conditions. **B**) Left, exemplar traces showing Ca^2+^ signals (indicated by Fura-2 fluorescence ratio) triggered in DRG neurones from NMR by Angiotensin II (200nM) II (Ang-II) alone or together with Pyr6 (2μM) in Ca^2+^-free extracellular media followed by subsequent Ca^2+^ add back. Right, Right, summary data obtained from Ca^2+^ imaging experiments shown in left, showing peak Ca^2+^ entry (SOCE) observed in DRG neurones from NMR under different conditions. Data are mean ± sem from ≥60 DRG neurons (per each experimental condition) plated on 3 petri dishes, each imaged on different days. Averages were compared using Student’s t test, 2-tailed, unpaired; p<0.05 was considered to be significant.

## CONCLUSION

In this study, we functionally and pharmacologically characterised SOCE in non-excitable (skin fibroblasts) and excitable (DRG neurons) NMR cells, finding that NMR cells exhibit robust SOCE which was sensitive to some widely-used selective modulators of this pathway, similar to what is observed in human and other mammalian cells. Considering the key role that SOCE plays in a multitude of cellular processes, from refilling depleted intracellular Ca^2+^ stores to gene transcription, cytokine secretion and metabolism[3], it is perhaps not surprising that SOCE activity appears normal in NMR cells, but this was still worthy of investigation considering the unusual physiology of NMRs[6]. What remains to be determined is whether the SOCE measured was reliant on Orai1 and Stim1 proteins as is the case in other mammalian cell types, but the expression of Orai1 and Stim1 in NMR SV40 fibroblasts indicates that they are likely involved. In addition, the potential contributions and physiological roles of Stim2 and Orai 2 and Orai3 also need to be investigated. Future studies may also include cloning and functional characterisation of individual Stim and Orai isoforms, determining their fundamental biophysical properties and sensitivity to pharmacological modulators.

## ACKNOWLEDGEMENT

Z.Z.H. was supported by a Cambridge Trust Cambridge International Scholarship.

## AUTHOR CONTRIBUTIONS

TR conceived the project and discussed with ESS regarding experimental planning. PD performed all the Ca^2+^ imaging studies; PUM, LQ and ESS provided NMR SV40 fibroblasts and NMR-DRG neurons; ZZH helped in confocal microscopy. TR wrote the manuscript with contributions from all. TR overall supervised the project, with regular consultation with ESS.

---

**Supplementary Figure 1**

**Figure S1. Alignment of full length Orai1 sequences of the naked mole rat (nmr), human, rat and mouse**. ClustalOmega in its default setting was used to perform multiple sequence alignment of the amino acid sequences corresponding mouse_Orai1 (Uniprot id: Q8BWG9), rat_Orai1 (Uniprot id: Q5M848), nmr_Orai1 (NCBI accession no. XP_004843916.1) and human_Orai1 (Uniprot id: Q96D31). The calculated Percent Identity Matrices between sequence pairs were >87.

**Supplementary Figure 2**

**Figure S2. Alignment of full length Orai1 sequences of the naked mole rat (nmr), human, rat and mouse**. ClustalOmega in its default setting was used to perform multiple sequence alignment of the amino acid sequences corresponding nmr_Orai2 (Uniprot id: G5AXA7), human_Orai2 (Uniprot id: Q96SN7), mouse_Orai2 (Uniprot id: Q8BH10) and rat_Orai2 (Uniprot id: B0BNI3). The calculated Percent Identity Matrices between sequence pairs were >95.

**Supplementary Figure 3**

**Figure S3. Alignment of full length Orai3 sequences of the naked mole rat (nmr), human, rat and mouse**. ClustalOmega in its default setting was used to perform multiple sequence alignment of the amino acid sequences corresponding mouse_Orai3 (Uniprot id: Q6P8G8), rat_Orai3 (Uniprot id: Q6AXR8), nmr_Orai3 (NCBI accession no. XP_004856256.1) and human_Orai3 (Uniprot id: Q9BRQ5). The calculated Percent Identity Matrices between sequence pairs were >87.

**Supplementary Figure 4. Alignment of full length Stim1 sequences of the naked mole rat (nmr), human, rat and mouse**. ClustalOmega in its default setting was used to perform multiple sequence alignment of the amino acid sequences corresponding nmr_Stim1 (Uniprot id: A0A0P6JBY0), mouse_Stim1 (Uniprot id: P70302) and rat_Stim1 (Uniprot id: P84903). The calculated Percent Identity Matrices between sequence pairs were >94.

**Supplementary Figure 5**

**Figure S5. Alignment of full length Stim2 sequences of the naked mole rat (nmr), human, rat and mouse**. ClustalOmega in its default setting was used to perform multiple sequence alignment of the amino acid sequences corresponding mouse_Stim2 (Uniprot id: P83093), rat_Stim2 (Uniprot id: A0A8I6AK33), nmr_Stim2 (Uniprot id: G5CB19) and human_Stim2 (Uniprot id: Q9P246). The calculated Percent Identity Matrices between sequence pairs were >93.

**Supplementary Figure 6**

**Figure S6. Validation of FITC-conjugated antibodies specific for hOrai1 and hStim1 in Jurkat T cells**. Exemplar confocal images showing expression of endogenous Orai and Stim proteins in Jurkat T cells that were fixed and stained with FITC-conjugated antibodies raised against hOrai1 and Stim1; nuclei were stained with DAPI and shown in blue. Scale bar, 50μm.

