## Supplementary figures and images for "Functional characterization of the store-operated calcium entry pathway in naked mole-rat cells"

### Supplementary Fig.1

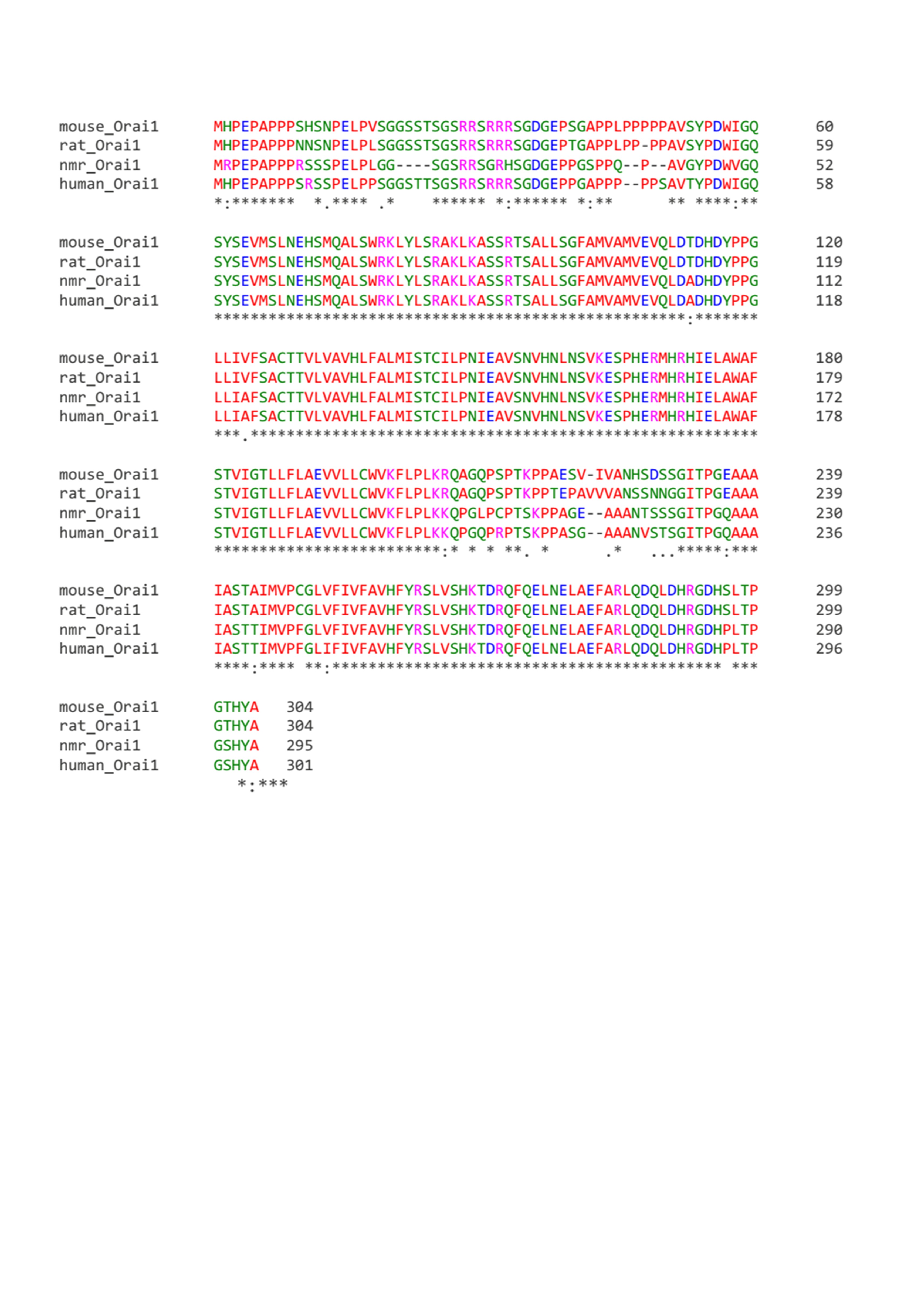

### Supplementary Fig.2

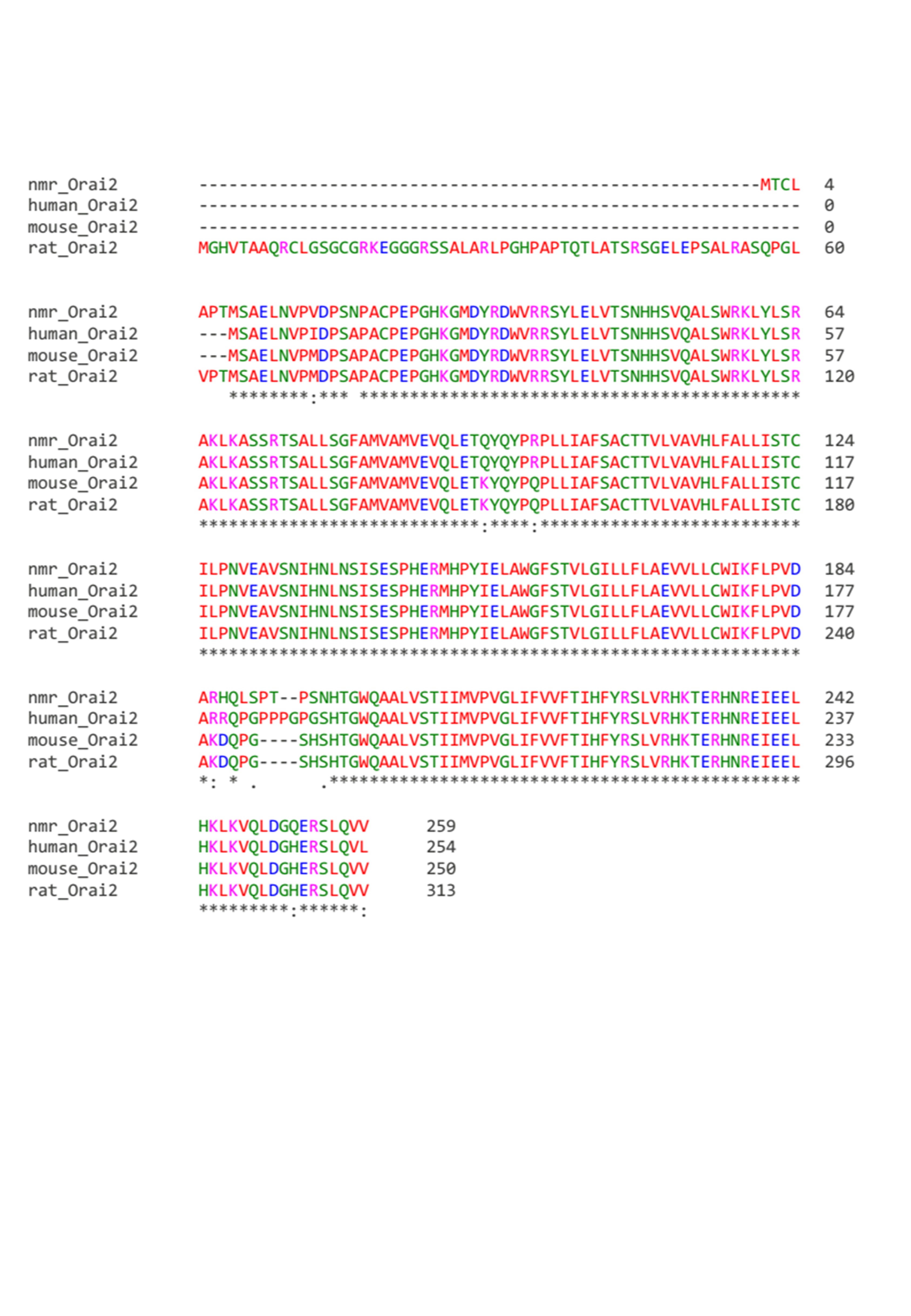

### Supplementary Fig.3

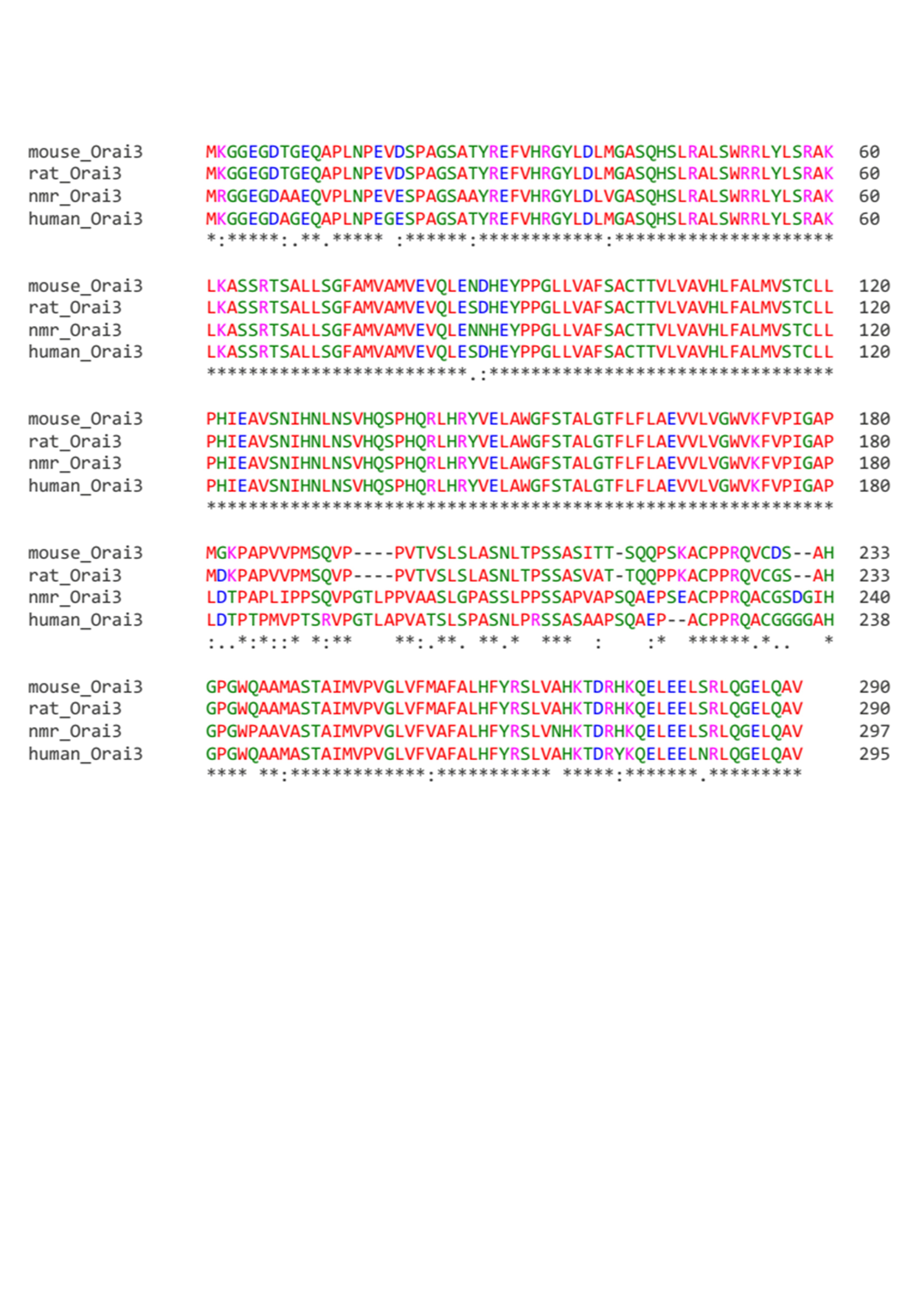

### Supplementary Fig.4

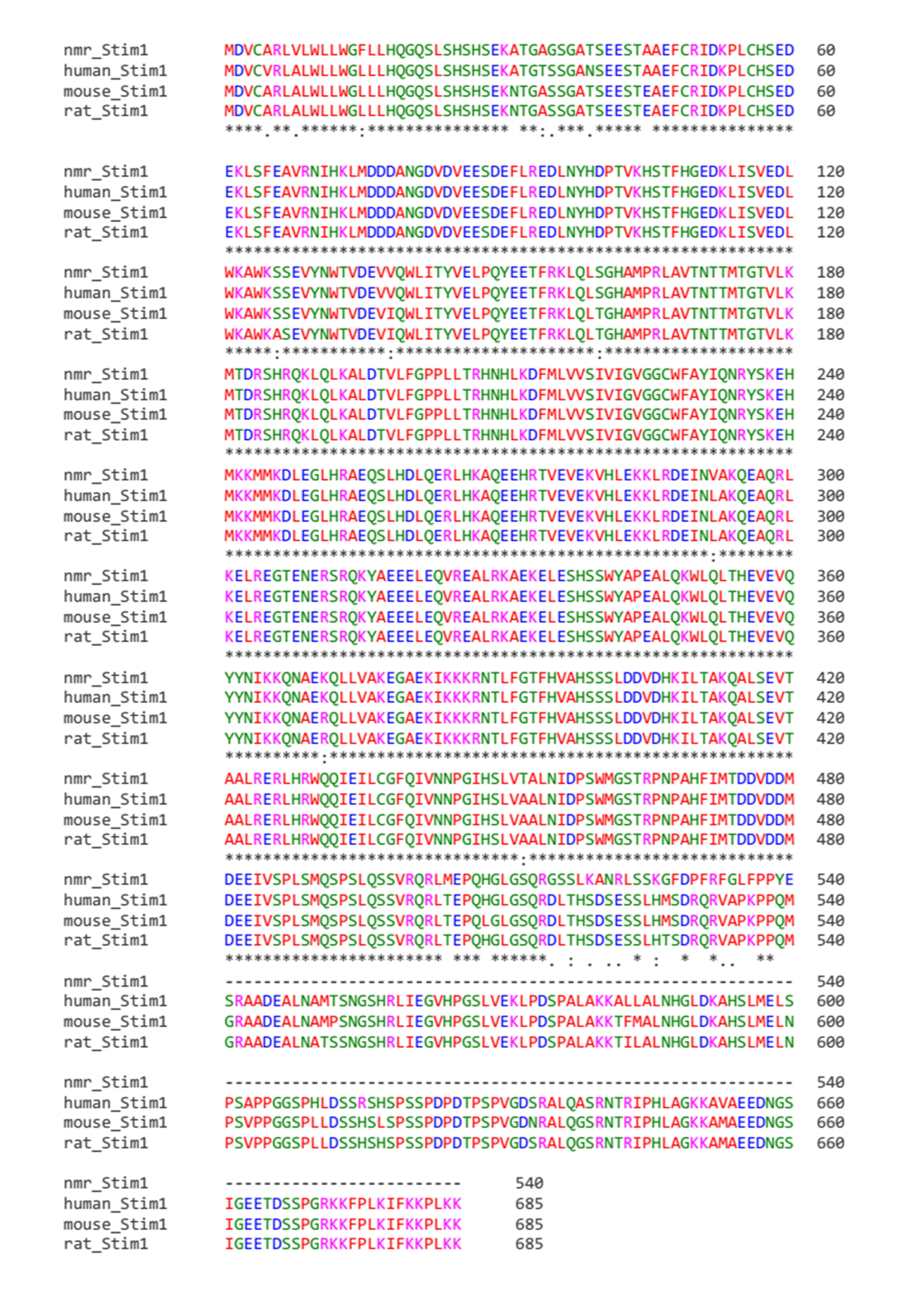

### Supplementary Fig.5

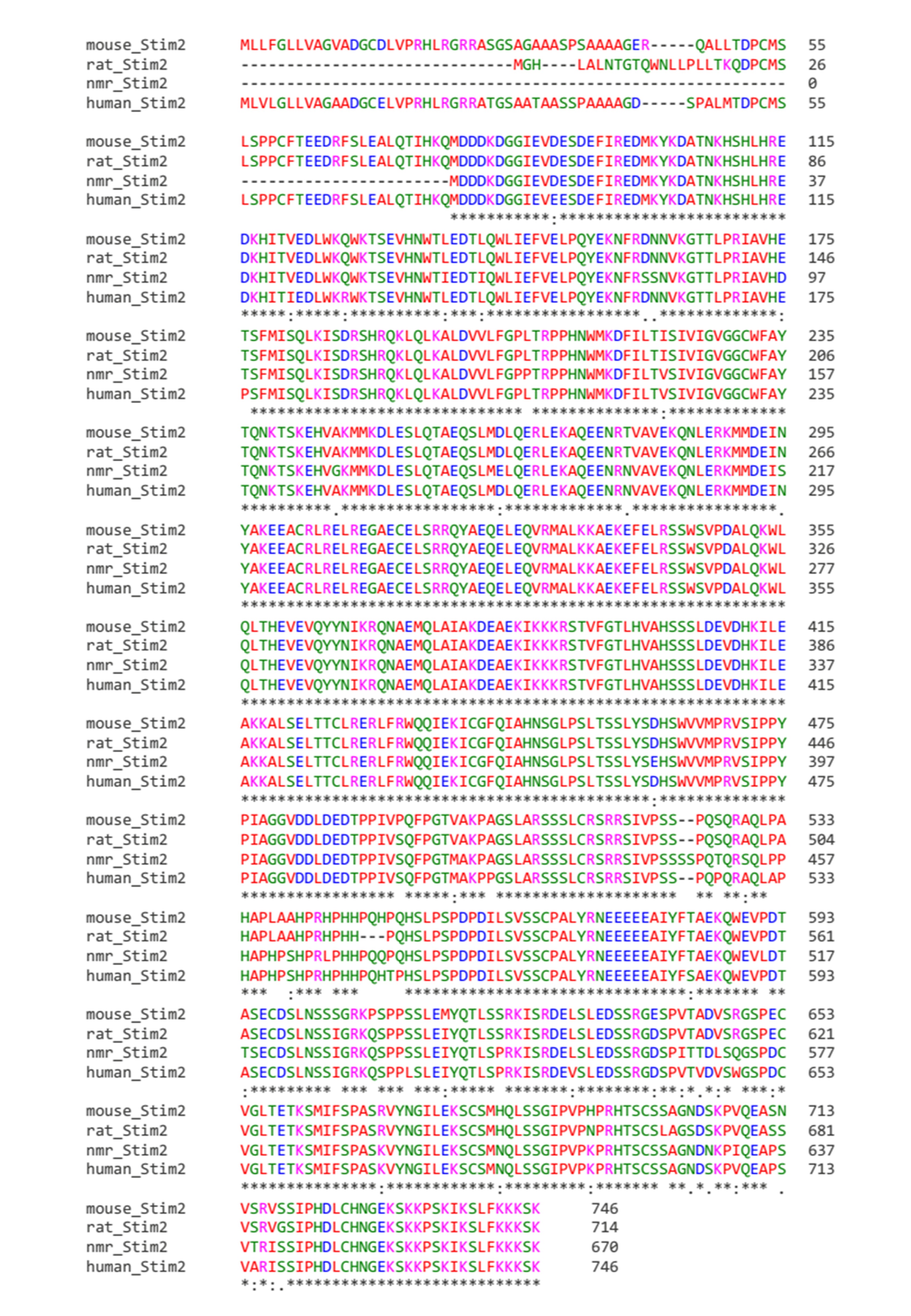

### Supplementary Fig.6

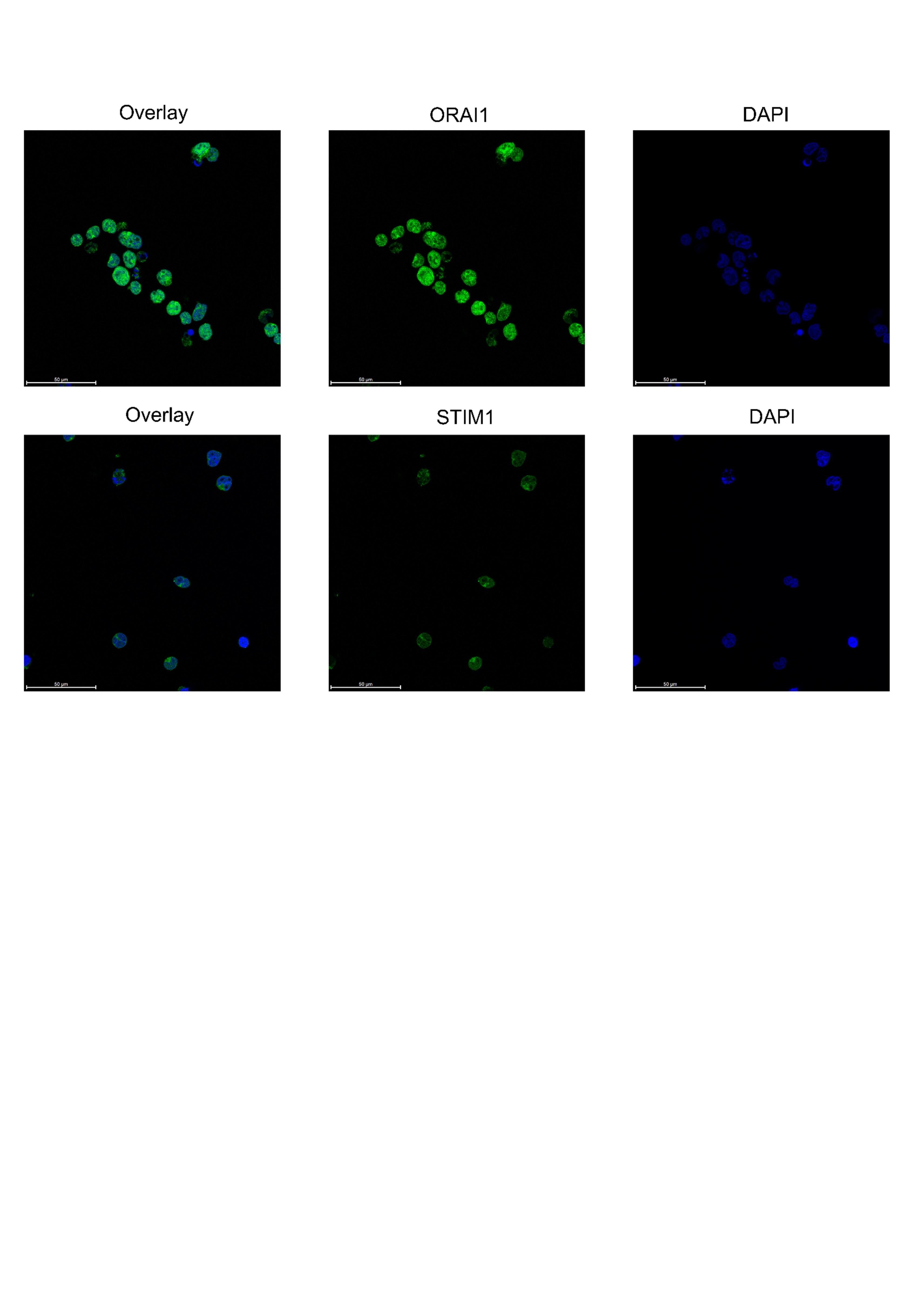
